# Bacterial adaptation to diet is a key evolutionary force shaping *Drosophila-Lactobacillus* symbiosis

**DOI:** 10.1101/222364

**Authors:** Maria Elena Martino, Pauline Joncour, Ryan Leenay, Hugo Gervais, Malay Shah, Sandrine Hughes, Benjamin Gillet, Chase Beisel, François Leulier

## Abstract

Animal-microbe facultative symbioses play a fundamental role in ecosystem and organismal health (1–3). Yet, due to the flexible nature of their association, the selection pressures acting on animals and their facultative symbionts remain elusive (4, 5). Here, by applying experimental evolution to a well-established model of facultative symbiosis: *Drosophila melanogaster* associated with *Lactobacillus plantarum*, one of its growth promoting symbiont (6, 7), we show that the diet, instead of the host, is a predominant driving force in the evolution of this symbiosis and identify the mechanism resulting from the bacterial adaptation to the diet, which confers host growth benefits. Our study reveals that adaptation to the diet can be the foremost step in the determination of the evolutionary course of a facultative symbiosis.

## Main Text

In facultative symbioses, microbes do not persistently colonize the host; nevertheless, they confer essential benefits to their animal partners (*8*, *9*). The flexible nature of these relationships suggests that there are reciprocal costs and benefits associated with maintaining such symbiosis (*3*, *9*, *10*). However, the ecological and evolutionary forces that drive the emergence and evolution of the benefits that facultative symbionts confer to their animal hosts remain largely elusive. To address this question, we experimentally tested microbial evolution using *Drosophila melanogaster* associated with one of its most abundant facultative symbionts, *Lactobacillus plantarum*, with whom it establishes nutritional mutualism (*9*, *11*–*14*). As growth promotion during undernutrition is one of the major advantages conferred by *L. plantarum* to its animal host (*11*, *15*), we asked if and how this bacterium can increase its potential to support animal growth while evolving with its host. To this end, we performed experimental evolution of NIZO2877 (*Lp*^NIZO2877^), a strain of *L. plantarum* isolated from processed human food (*16*), which was previously shown to moderately promote growth both in *Drosophila* and mice (*11*, *15*). We mono-associated germ-free (GF) *Drosophila* eggs with a fully sequenced clonal population of *Lp*^NIZO2877^ on a low-nutritional diet and studied the partners for 20 *Drosophila* generations (i.e 313 days, which correspond to about 2000 bacterial generations; see Methods and Fig. S1-2). At each generation, we selected the first emerging pupae carrying a subpopulation of *L. plantarum* strains, and transferred them to a new sterile diet (Methods; Fig. S1). The adults rapidly emerged from the pupae and deposited the new embryos and their associated *L. plantarum* strains that subsequently colonized and propagated in the new environment. We then isolated the *Lp*^NIZO2877^-evolved strains associated with the adult flies eclosed from the transferred pupae, selected a representative set of isolates and measured individually their growth promoting capacity on an independent set of GF fly larvae. After only two fly generations (i.e. after about 124 bacterial generations, Fig. 1A,B), we identified a few evolved *Lp*^NIZO2877^ strains that significantly improved larval growth and accelerated pupariation compared to the ancestor strain. Specifically, the evolved strains exhibited the same effect as *Lp*^WJL^, a potent *L. plantarum* growth promoting strain (*15*) (Fig. 1A,B). These results show that the evolution of *Lp*^NIZO2877^ in the context of its symbiosis with *Drosophila* leads to the rapid improvement of *L. plantarum* animal growth promotion (Fig. S3).

**Fig. 1.**
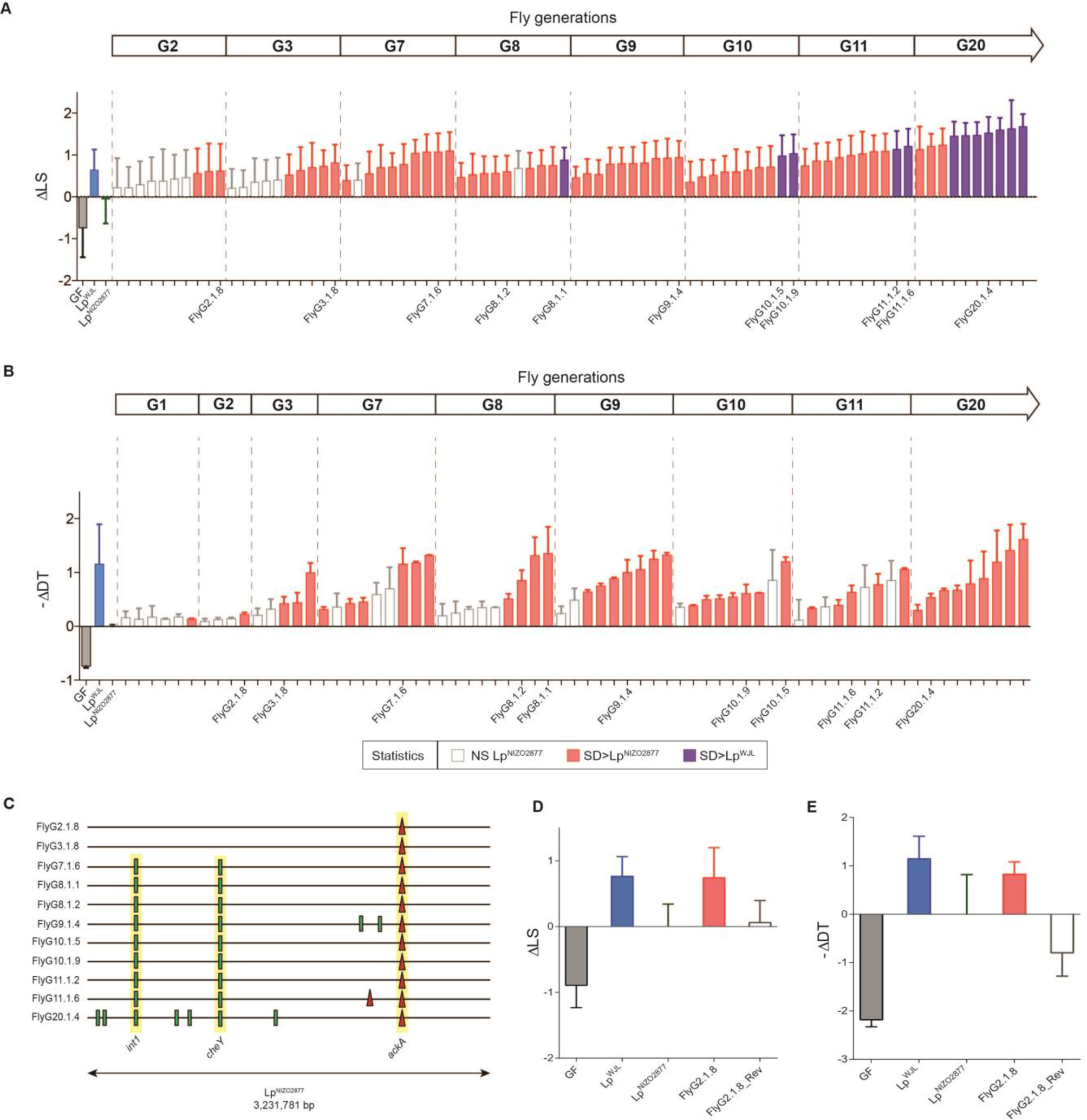
Experimental evolution of *L. plantarum* with *Drosophila melanogaster* improves its growth promoting effect. **A,** Longitudinal size of larvae (LS) measured 7 days after egg deposition (AED) on poor nutrient diet. Larvae were kept germ-free (GF) or associated with *Lp*^NIZO2877^ (ancestor), *Lp*^WJL^ (growth-promoting *L. plantarum* strain) or *Lp*^NIZO2877^-evolved strains. The delta (ΔLS) between the size of larvae associated with the respective condition and the size of larvae associated with *Lp*^NIZO2877^ is shown from *Drosophila* generation 2 (G2) to generation 20 (G20). *Lp*^NIZO2877^-evolved strains that exhibited a significant difference at promoting larval growth compared to their ancestor (Student’s t test: p<0.05) are shown in red. *Lp*^NIZO2877^-evolved strains that exhibited a significant difference at promoting larval growth compared to the beneficial *L. plantarum Lp*^WJL^ strain are shown in purple. **B,** Developmental timing (DT) of individuals that were kept GF or associated with *Lp*^NIZO2877^, *Lp*^WJL^ or *Lp*^NIZO2877^-evolved strains isolated from *Drosophila* G1 to G20. The minus delta (-ΔDT) between the mean time of emergence of 50% of the pupae associated with the respective condition and the mean time of emergence of 50% of the pupae associated with *Lp*^NIZO2877^ is shown in the graph. *Lp*^NIZO2877^-evolved strains that exhibited a significant difference at accelerating developmental timing compared to the ancestor (Student’s t test: p<0.05) are shown in red. The evolved strains that have been selected for further analyses are labelled on the x axis. **C,** Mutations identified in *Lp*^NIZO2877^-evolved strains from *Drosophila* generation 2 (G2) to generation 20 (G20) represented along *Lp*^NIZO2877^ genome. The genome of each evolved strain is represented as a horizontal line. Red triangles indicate deletions and small green bars show single nucleotide polymorphisms. Mutations occurring in the same gene of different strains and fixed along the experimental evolution are highlighted in yellow (*int1*, *cheY*, *ackA*). **D,** Longitudinal size of larvae measured 7 days AED on poor nutrient diet. Larvae were kept germ-free (GF) or associated with *Lp*^NIZO2877^*, Lp*^WJL^, FlyG2.1.8 or with FlyG2.1.8-reverted strain (FlyG2.1.8^Rev^). The delta (ΔLS) between the size of larvae associated with the respective *L. plantarum* strain and the size of larvae associated with *Lp*^NIZO2877^ is shown. **E,** Developmental timing (DT) of individuals that were kept GF or associated with *Lp*^NIZO2877^*, Lp*^WJL^, FlyG2.1.8 or with FlyG2.1.8^Rev^ strain. The minus delta (-ΔDT) between the mean time of emergence of 50% of the pupae associated with the respective condition and the mean time of emergence of 50% of the pupae associated with *Lp*^NIZO2877^ is shown in the graph.

To identify the genetic changes underlying the rapid microbial adaptation responsible for the improved growth of the host, we sequenced the genomes of 11 evolved *Lp*^NIZO2877^ strains (Table S1, replicate 1) with increased host growth promoting potential across the 20 *Drosophila* generations. We identified a total of 11 mutations, including nine single-nucleotide polymorphisms (SNPs) and two small deletions (Fig. 1C; Table S2). In particular, in the strain isolated from the second fly generation (FlyG2.1.8), we found a single change in the genome within one of the three acetate kinase genes (*ackA*). Remarkably, this first mutation was subsequently fixed and strictly correlated with the improved animal growth phenotype (Fig. 1C).

To test the repeatability of this finding, we conducted an independent replicate of *L. plantarum* experimental evolution while in symbiosis with *Drosophila*. Both the phenotypic and genomic evolution of *L. plantarum* were again obtained: *Lp*^NIZO2877^ improved its animal growth promoting potential by rapidly acquiring and fixing mutations, including variants in the *ackA* gene (Fig. S4, Table S2). In the first experiment, the evolved *Lp*^NIZO2877^ strains with improved animal growth potential all carried a three-nucleotide deletion in the *ackA* gene that removed one proline residue. From the second replicate, the evolved strains carried a SNP that resulted in a premature stop-codon leading to protein truncation (Fig. S5). These independently isolated mutations likely generate an inactive *ackA* protein. Following the fixation of *ackA* variants, additional mutations appeared in both replicates of *L. plantarum* experimental evolution, which seem to further improve its symbiotic benefit (Fig. 1A, Fig. S4A). Nevertheless, the two evolved strains each bearing only one mutation in *ackA* (FlyG2.1.8 and FlyG3.1.8) already showed a statistically significant *Drosophila* growth improvement compared to their ancestor (Fig. 1A,B). Based on these observations, we propose that the *de novo* appearance of the *ackA* mutation is the first fundamental step in shaping the evolutionary trajectory in the *Lp*^NIZO2877^/*Drosophila* symbiosis model.

To fully establish that *ackA* mutation is responsible for the evolution of *Lp*^NIZO2877^/*Drosophila* symbiosis, we employed CRISPR-Cas9 to re-insert the deleted CCT triplet in the FlyG2.1.8 *ackA* locus (Methods; Fig. S6), so that we genetically revert the *ackA* allele in the FlyG2.1.8 isolate back to its ancestral form (*17*). The reverted strain (FlyG2.1.8^Rev^) bearing the ancestral *ackA* allele lost its increased capacity to promote animal growth when compared to the ancestor strain (Fig. 1D,E). These results therefore demonstrate that the *ackA* mutation in *Lp*^NIZO2877^ is a causative change resulting in faster and increased *Drosophila* growth.

To investigate the complete *L. plantarum* population dynamics during *Drosophila* symbiosis evolution, we sequenced the metagenome of whole bacterial population samples across the 20 *Drosophila* generations of the first replicate experiment. We identified both segregating and fixed mutations and tracked their frequencies through time (Methods). We found that the *ackA* mutation was the first variant to appear in the population. Remarkably, the *ackA* variant showed a rapid selective sweep and became fixed as early as after three *Drosophila* generations (Fig. 2A). This observation suggests a competitive advantage of the evolved *Lp*^NIZO2877^ strains bearing this variant. To test this hypothesis, we performed a competition assay between the ancestral *Lp*^NIZO2877^ strain and the derived FlyG2.1.8 isolate in symbiosis with *Drosophila* (Methods, Fig.2B, Fig. S7). We find that the evolved strain bearing only the *ackA* mutation starts outcompeting the ancestor strain as early as after one day, demonstrating that the *ackA* mutation confers a strong competitive advantage in symbiosis with *Drosophila*. To test whether such advantage requires the host’s presence, we performed the same competition assay by inoculating only the bacterial strains on the *Drosophila* nutritional environment (i.e. the diet). Surprisingly, we observed that FlyG2.1.8 outcompeted the ancestral strain even when the *Drosophila* host is absent (Fig. 2C). Therefore, the competitive advantage of *L. plantarum* isolates bearing the *ackA* variant is likely independent of the animal host.

**Fig. 2.**
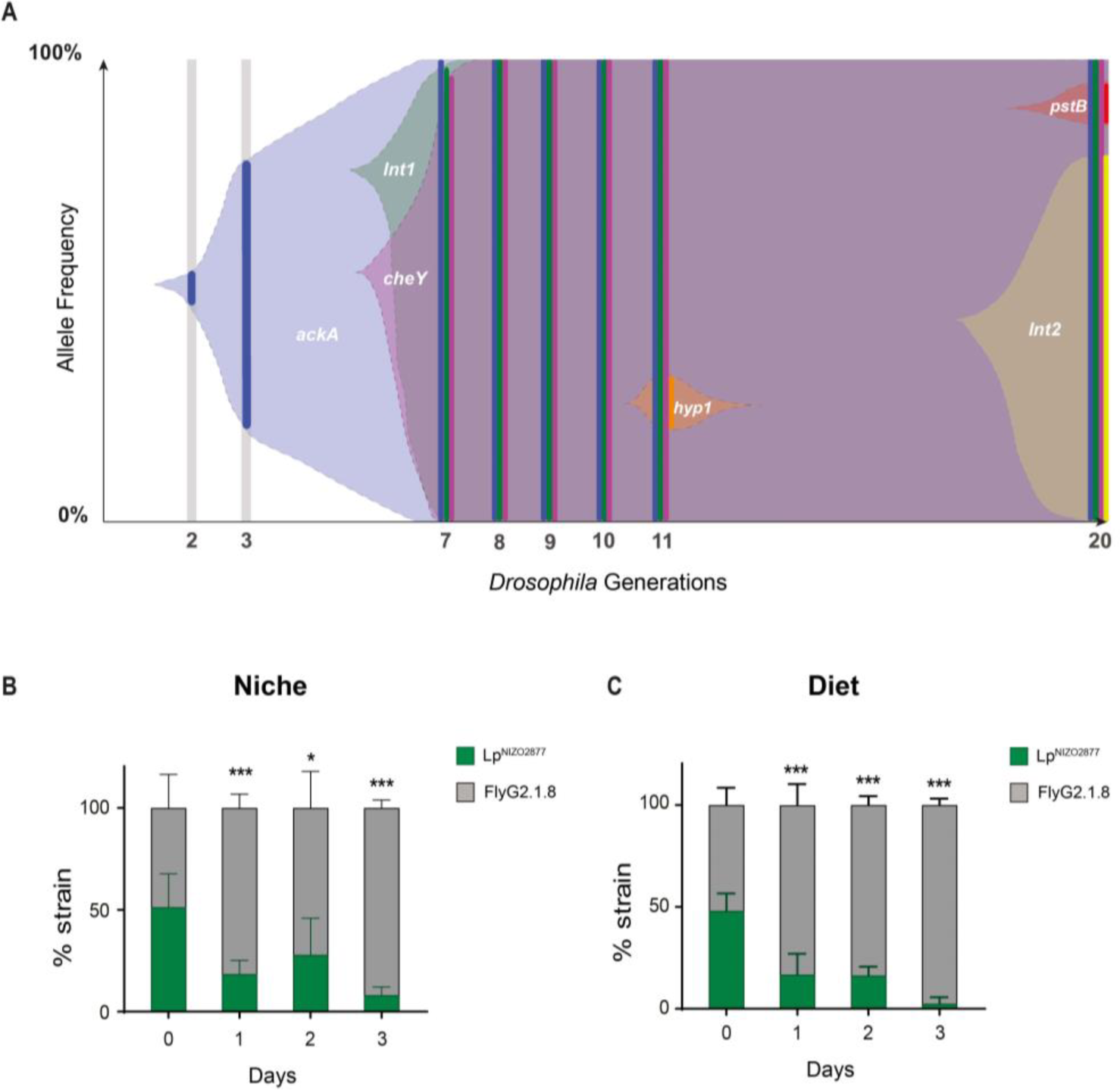
*Lp*^NIZO2877^-evolved strain shows higher fitness compared to the ancestor. **A,** Muller diagram showing the genome evolutionary dynamics of *Lp*^NIZO2877^ population (I replicate) along 20 *Drosophila* generations. The y-axis shows the percentage of the detected frequencies of each mutation (plain colours). Shaded areas represent the inferred mutation frequencies. Lower axis shows the fly generation where the sampling took place. **B-C,** 1:1 competitive assay between *Lp*^NIZO2877^ and *Lp*^NIZO2877^*-*evolved strain (FlyG2.1.8) in poor nutrient diet with *Drosophila* larvae (**B**) and without *Drosophila* larvae (**C**). Bars represent the percentage of each strain detected in each sample (Niche or Diet) by qPCR. \**P* < 0.05, \*\*\**P* < 0.01, obtained by Student’s t-test.

Intrigued by this result, we questioned whether the animal host has an influence on the evolution of its symbiotic bacteria. To test this, we experimentally evolved *Lp*^NIZO2877^ in the same low-yeast fly diet, but without *Drosophila* (Methods; Fig. S8) and tested their capacity to promote fly growth throughout the course of the experimental evolution. Strikingly, in two parallel experiments, the *Lp*^NIZO2877^ strains evolved in the absence of the host also increased their ability to promote *Drosophila* growth (Fig. 3A,B). Furthermore, genome sequencing of single evolved isolates from both experiments again revealed the acquisition of novel mutations in the *ackA* gene (Fig. 3C; Fig. S9). Taken together, these findings show that the genomic evolution of *L. plantarum* is driven by the adaptation to host nutritional environment, rather than to its host *per se*; the acquisition of the *ackA* variant is sufficient to drive the adaptive process to the nutrition, which ultimately results in the improvement of *L. plantarum* symbiotic effect on *Drosophila*.

**Fig. 3.**
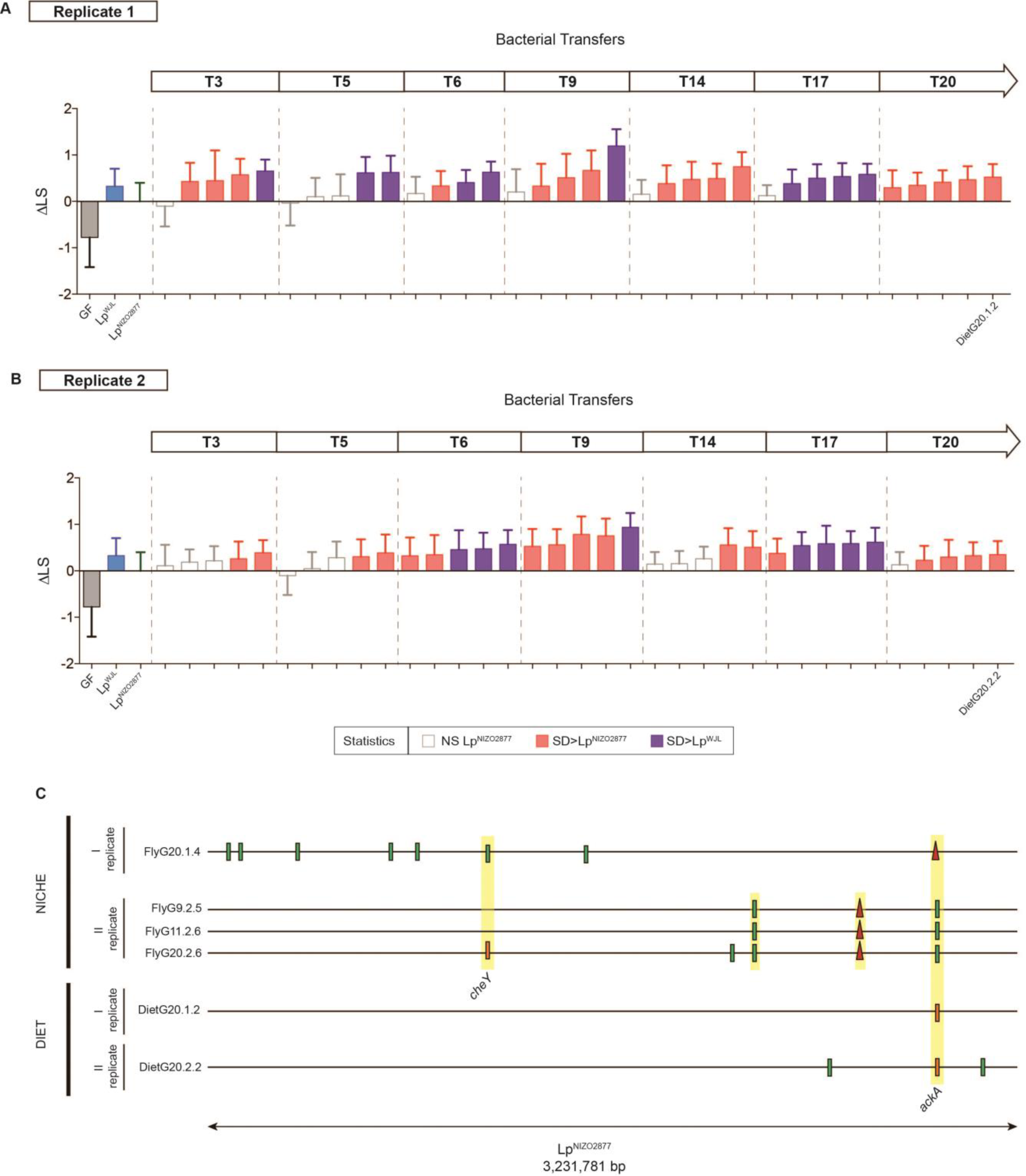
*Lp*^NIZO2877^ adaptation to the diet increases its host’s growth. **A,B,** Longitudinal size of larvae (LS) measured 7 days after egg deposition (AED) on poor nutrient diet. Larvae were kept germ-free (GF) or associated with *Lp*^NIZO2877^*, Lp*^WJL^ and with *Lp*^NIZO2877^-evolved strains evolved in poor nutrient diet in the absence of *Drosophila*. The delta (ΔLS) between the size of larvae associated with *Lp*^NIZO2877^-evolved strains and the size of larvae associated with *Lp*^NIZO2877^ is shown from transfer 3 (T3) to transfer 20 (T20) for the first replicate (**A**) and the second replicate (**B**) of evolution. *Lp*^NIZO2877^-evolved strains that exhibited a significant difference at promoting larval growth compared to their ancestor (Student’s t test: p<0.05) are shown in red. *Lp*^NIZO2877-^evolved strains that exhibited a significant difference at promoting larval growth compared to the beneficial *L. plantarum Lp^WJ^*L strain are shown in purple. The evolved strains that have been selected for further analyses are labelled on the x axis. **c,** Mutations identified in *Lp*^NIZO2877^-derived strains of all replicates evolved in poor nutrient diet with *Drosophila* larvae (Niche) and in poor nutrient diet without *Drosophila* larvae (Diet). Each evolved strain genome is represented as a horizontal line. Red triangles indicate deletions and small bars shows single nucleotide polymorphisms. Different colours indicate different variants. Mutations occurring in the same gene and fixed along the experimental evolution are highlighted in yellow. The genes mutated in independent replicates of experimental evolution are labelled (*cheY*, *ackA*).

We next investigated how *L. plantarum* adaptation to the nutritional environment enhances *Drosophila* growth. We postulated that *L. plantarum* adaptation to the specific nutritional environment of *Drosophila* would lead to the production of metabolites that are beneficial for *Drosophila* growth. To test this hypothesis, we analyzed the metabolome of *Drosophila* diets colonized with either *Lp*^NIZO2877^ or the evolved FlyG2.1.8 strain that bears only the *ackA* variant. Among all of the metabolites differentially detected in the substrate (Table S6), we observed a significant and robust increase in the levels of N-acetyl-amino-acids in the diet processed by the evolved strain (Fig. 4A). Specifically, N-acetyl-glutamine is one of the most differentially represented compounds between the two conditions. We therefore tested whether N-acetylglutamine is sufficient to improve the animal growth promoting capacity of *Lp*^NIZO2877^. Remarkably, we find that, when N-acetyl-glutamine is added in a dose-dependent manner in the diet, the ancestor strain *Lp*^NIZO2877^ is able to recapitulate the beneficial effect conferred by FlyG2.1.8 on *Drosophila* growth (Fig. 4B). We then asked whether N-acetyl-glutamine enhances fly growth by improving *Lp*^NIZO2877^ fitness. To test this, we performed a competition assay between *Lp*^NIZO2877^ and FlyG2.1.8 strains in the host diet supplemented with 0.1g/L of N-acetyl-glutamine. We find that FlyG2.1.8 outcompetes the ancestor strain even in presence of N-acetyl-glutamine (Fig. S10). This result indicates that N-acetyl-glutamine does not confer a competitive advantage to *Lp*^NIZO2877^ over FlyG2.1.8 while growing on the diet; nevertheless it benefits the host physiology. Taken together, these findings establish N-acetyl-amino-acids, and in particular N-acetyl-glutamine, as molecules produced by the evolved *L. plantarum* strains during growth on the *Drosophila* diet, which enhance *Drosophila* growth but not *Lp*^NIZO2877^ fitness.

**Fig. 4.**
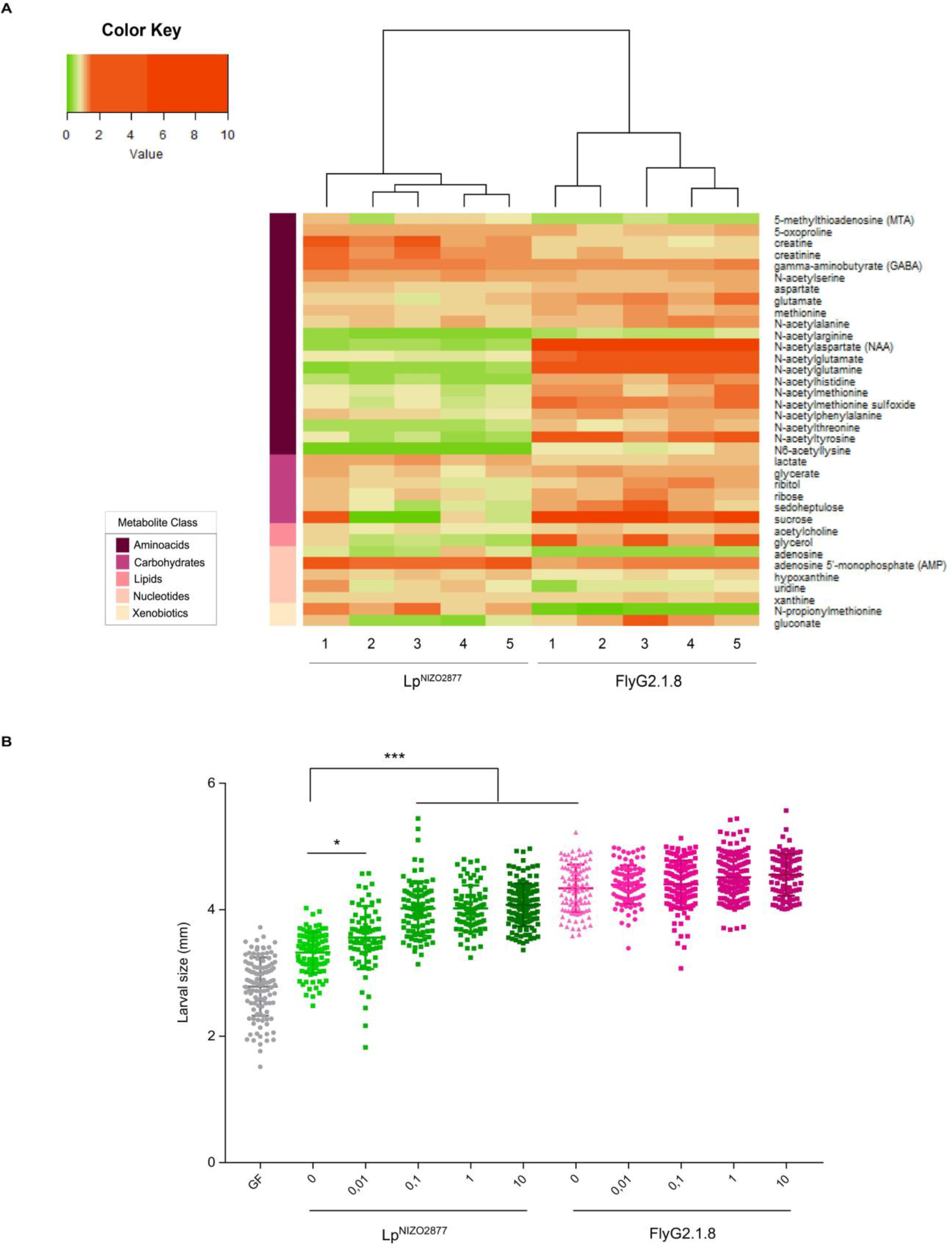
N-acetyl-glutamine recapitulates the beneficial effect of FlyG2.1.8 on *Lp*^NIZO2877-^associated larvae. **A,** Heat map showing the metabolites that differ significantly between experimental groups (*Lp*^NIZO2877^ and FlyG2.1.8) (two-sided *t*-tests p<0.05). The heat map was generated with *heatmap.2* function in R. The compounds are ordered by the metabolite class given by the left scale. **B,** Longitudinal size of larvae (n > 60 larvae/group) measured 7 days after egg deposition on poor nutrient diet supplemented with different concentrations (g/L) of N-acetyl-glutamine (x axis). Larvae were kept germ-free (no supplementation of N-acetylglutamine) or associated with *Lp*^NIZO2877^ (ancestor) and with Fly.G2.1.8 (evolved strain). Larval size is shown as mean ± s.e.m. * P < 0.05, *** P < 0.01.

Our results uncover the nature of an adaptive process of *L. plantarum* while in symbiosis with its fly host. To our knowledge, this is the first direct experimental evidence showing that the host nutritional environment, and not the host *per se,* drives microbial adaptation and metabolic changes that alter the functional outputs of a facultative nutritional symbiosis. In our experimental context, the dietary substrate asserts the predominant selective pressure dictating the evolutionary change of facultative symbiotic bacteria and their consequent benefits to host physiology. Rapid adaptation of *L. plantarum* to the host nutritional environment occurred in multiple independent experimental lineages through the parallel fixations of different variants of a single gene, the acetate kinase *ackA*. This is a spectacular case of parallel evolution, indicating that the *ackA* mutation is the preferred or possibly the unique means for *L. plantarum*^NIZO2877^ to adapt to its host nutritional environment. These harsh nutritional conditions of our experimental setting affect *L. plantarum* physiology by delaying its growth (Fig. S2). It was shown that the expression of *L. plantarum ackA* (*ack2* in the *L. plantarum* reference strain WCFS1) is down-regulated at low growth rates suggesting that silencing *ackA* would be required to cope with poor growth condition (*18*). This observation may explain the observed strong selection pressure on *ackA* in our experimental settings, which led to the rapid *de novo* emergence of different variants in the population (Fig. 2A). As a consequence, the strong competitive advantage given by these mutations led to their fixation (Fig. 2). Indeed, the *ackA* mutations found in the independent lineages of adaptive evolution improve the fitness of *L. plantarum* cells on the fly diet (Fig. S11), and leads to the accumulation of bacterial products, such as N-acetyl-glutamine, that enhance host growth. However, N-acetyl-glutamine does not per se improve bacterial fitness so it remains elusive how *ackA* variants confers competitive advantage to *L. plantarum* cells on the fly diet. Our results indicate that these mutations possibly cause a shift in the metabolism of *L. plantarum* by modifying the usage of cellular acetyl groups, which would confer benefits to *Drosophila* larvae growth. *ackA* participates in the reversible conversion of acetate to acetyl-phosphate; *ackA* variants might impede this reaction, and therefore shunt the pools of cellular acetyl groups into different metabolic routes leading to the accumulation of other acetylated compounds, such as N-acetyl-amino-acids, which, once secreted, are consumed and beneficial to the host. Our results identify *ackA* as the first target of selection exerted by the nutritional environment on *Lp*^NIZO2877^. Due to the high genetic variability of *L. plantarum* species (*19*), we posit that such target hinges upon the genomic background of *Lp*^NIZO2877^. According to their network of genetic polymorphisms, other non-beneficial isolates might mutate different genes in order to adapt to the host environment and improve their symbiotic benefit. Regardless of the specificity of selection target, our findings determine that the host nutritional environment is the first driving force of such evolution.

Understanding how evolutionary forces shape host-microbe symbiosis is essential to comprehend the mechanisms of their functional influence. Using the facultative nutritional mutualism between *Drosophila* and *Lactobacillus plantarum* as a model, our results reveal that the primary selection pressure acting on *Lactobacillus plantarum* originates from the nutritional substrate alone, which is strong enough to drive the rapid fixation of a *de novo* mutation. The resulting genetic change confers a fitness advantage to the evolved bacteria and triggers a metabolic adaptation in bacterial cells, which is quickly capitalized by *Drosophila* as a physiological growth advantage, and symbiosis can henceforth be perpetuated. Our results do not rule out the possibility that the animal host might exert additional selection pressure on its bacterial partners. Indeed, *Drosophila* is also known to directly impact the fitness of its own microbiota through the activity of innate immune effectors (*20*, *21*) or the secretion of bacterial maintenance factors (*22*). Nevertheless, our findings demonstrate the utmost importance of the shared nutritional substrate in the evolution of *Drosophila*-*L. plantarum* symbiosis.

Symbiosis is an evolutionary imperative and facultative symbioses are widespread in nature. Despite their unequivocal diversity, animal-microbe symbioses share striking similarities (*4*) and nutrition often plays a major role in shaping the composition of symbiotic microbial communities (*23*–*28*). Our results provide the first direct experimental evidence that nutrition drives the evolution of a bacterial symbiont and, given that other animal and microbe partners have likely faced nutritional challenges over time, common evolutionary trajectories might have occurred. We therefore posit that bacterial adaptation to the diet can be the first step in the emergence and perpetuation of facultative animal-microbe symbioses. Our work provides another angle to unravel the complex adaptive processes in the context of evolving symbiosis.

## Acknowledgements

We thank B. Prud’homme and colleagues at the Institut de Génomique Fonctionnelle de Lyon for critical reading of the manuscript. WCFS1 was a kind gift from Dr. Nikhil U. Nair. Plasmids pJP042 and pJP005 were both provided by the van-Pijkeren lab. pMSP3545 (CN#46888) and pCas9 (CN# 42876) were both obtained from Addgene. EC135 was provided to us by Tingyi Wen. We gratefully acknowledge support from the PSMN (Pôle Scientifique de Modélisation Numérique) of the ENS de Lyon for the computing resources. We thank University of Padua and Dr. Barbara Cardazzo for hosting M.E.M. during the last stages of this research.

## Author Contributions

M.E.M. and F.L. designed the project; M.E.M. and H.G. conducted the experiments; M.E.M. and P.J. conducted the bioinformatics analyses; R.L., M.S. and C.L.B. designed and performed the CRISPR-Cas9 engineering experiments; S.H. and B.G. generated the sequencing data; M.E.M. and F.L. analyzed the data and wrote the paper.

## Data and materials availability

All data is available in the main text or the supplementary materials.

## References

1. M. McFall-Ngai et al., Animals in a bacterial world, a new imperative for the life sciences. Proc. Natl. Acad. Sci. 110, 3229–3236 (2013).

2. F. Leulier et al., Integrative Physiology: At the Crossroads of Nutrition, Microbiota, Animal Physiology, and Human Health. Cell Metab. 25 (2017), pp. 522–534.

3. R. M. Fisher, L. M. Henry, C. K. Cornwallis, E. T. Kiers, S. A. West, The evolution of host-symbiont dependence. Nat. Commun. 8 (2017), doi:10.1038/ncomms15973.

4. K. R. Foster, J. Schluter, K. Z. Coyte, S. Rakoff-Nahoum, The evolution of the host microbiome as an ecosystem on a leash. Nature. 548 (2017), pp. 43–51.

5. A. E. Douglas, J. H. Werren, Holes in the hologenome: Why host-microbe symbioses are not holobionts. MBio. 7 (2016), doi:10.1128/mBio.02099-15.

6. R. C. Matos, F. Leulier, Lactobacilli-Host mutualism: “learning on the fly.” Microb. Cell Fact. 13, S6 (2014).

7. B. Erkosar, G. Storelli, A. Defaye, F. Leulier, Host-intestinal microbiota mutualism: “learning on the fly.” Cell Host Microbe. 13 (2013), pp. 8–14.

8. J. Ferrari, F. Vavre, Bacterial symbionts in insects or the story of communities affecting communities. Philos. Trans. R. Soc. B Biol. Sci. 366, 1389–1400 (2011).

9. A. E. Douglas, Lessons from studying insect symbioses. Cell Host Microbe. 10, 359–367 (2011).

10. P. Engel, N. A. Moran, The gut microbiota of insects - diversity in structure and function. FEMS Microbiol. Rev. 37 (2013), pp. 699–735.

11. G. Storelli et al., Lactobacillus plantarum promotes Drosophila systemic growth by modulating hormonal signals through TOR-dependent nutrient sensing. Cell Metab. 14, 403–14 (2011).

12. B. Erkosar et al., Pathogen Virulence Impedes Mutualist-Mediated Enhancement of Host Juvenile Growth via Inhibition of Protein Digestion. Cell Host Microbe. 18, 445–455 (2015).

13. D. Ma, G. Storelli, M. L. Mitchell, F. Leulier, Studying Host-Microbiota Mutualism in Drosophila: Harnessing the Power of Gnotobiotic Flies. Biomed. J., 285–293 (2015).

14. R. C. Matos et al., D-Alanylation of teichoic acids contributes to Lactobacillus plantarum-mediated Drosophila growth during chronic undernutrition. Nat. Microbiol. 2, 1635–1647 (2017).

15. M. Schwarzer et al., Lactobacillus plantarum strain maintains growth of infant mice during chronic undernutrition. Science. 351, 854–857 (2016).

16. M. E. Martino et al., Nearly complete genome sequence of Lactobacillus plantarum strain NIZO2877. Genome Announc. (2015), doi:10.1128/genomeA.01370-15.

17. W. Jiang, D. Bikard, D. Cox, F. Zhang, L. A. Marraffini, RNA-guided editing of bacterial genomes using CRISPR-Cas systems. Nat. Biotechnol. 31, 233–239 (2013).

18. P. Goffin et al., Understanding the physiology of Lactobacillus plantarum at zero growth. Mol. Syst. Biol. 6 (2010), doi:10.1038/msb.2010.67.

19. M. E. Martino et al., Nomadic lifestyle of Lactobacillus plantarum revealed by comparative genomics of 54 strains isolated from different habitats. Environ. Microbiol. 18 (2016), doi:10.1111/1462-2920.13455.

20. J. H. Ryu et al., Innate immune homeostasis by the homeobox gene Caudal and commensal-gut mutualism in Drosophila. Science (80-. ). 319, 777–782 (2008).

21. L. Guo, J. Karpac, S. L. Tran, H. Jasper, PGRP-SC2 Promotes Gut Immune Homeostasis to Limit Commensal Dysbiosis and Extend Lifespan. Cell. 156, 109–22 (2014).

22. Storelli et al., *Drosophila* Perpetuates Nutritional Mutualism by Promoting the Fitness of Its Intestinal Symbiont *Lactobacillus plantarum*. Cell Metab. (2018), doi:10.1016/j.cmet.2017.11.011

23. M. Groussin et al., Unraveling the processes shaping mammalian gut microbiomes over evolutionary time. Nat. Commun. 8 (2017), doi:10.1038/ncomms14319.

24. M. A. Conlon, A. R. Bird, The impact of diet and lifestyle on gut microbiota and human health. Nutrients. 7, 17–44 (2015).

25. C. A. Lozupone, J. I. Stombaugh, J. I. Gordon, J. K. Jansson, R. Knight, Diversity, stability and resilience of the human gut microbiota. Nature. 489, 220–230 (2012).

26. L. A. David et al., Host lifestyle affects human microbiota on daily timescales. Genome Biol. 15 (2015), doi:10.1186/gb-2014-15-7-r89.

27. B. D. Muegge et al., Diet drives convergence in gut microbiome functions across mammalian phylogeny and within humans. Science (80-. ). 332, 970–974 (2011).

28. S. Hacquard et al., Microbiota and host nutrition across plant and animal kingdoms. Cell Host Microbe. 17, 603–616 (2015).

29. L. K. Poulsen, T. R. Licht, C. Rang, K. A. Krogfelt, S. Molin, Physiological state of Escherichia coli BJ4 growing in the large intestines of streptomycin-treated mice. J. Bacteriol. 177, 5840–5845 (1995).

30. C. D. Packey et al., Molecular detection of bacterial contamination in gnotobiotic rodent units. Gut Microbes. 4, 361–370 (2013).

31. F. Widdel, Theory and measurement of bacterial growth. Di dalam Grundpraktikum Mikrobiol., 1–11 (2007).

32. C. a Schneider, W. S. Rasband, K. W. Eliceiri, NIH Image to ImageJ: 25 years of image analysis. Nat. Methods. 9, 671–675 (2012).

33. D. E. Deatherage, J. E. Barrick, Identification of mutations in laboratory-evolved microbes from next-generation sequencing data using breseq. Methods Mol. Biol. 1151, 165–188 (2014).

34. Y. Choi, G. E. Sims, S. Murphy, J. R. Miller, A. P. Chan, Predicting the Functional Effect of Amino Acid Substitutions and Indels. PLoS One. 7 (2012), doi:10.1371/journal.pone.0046688.

35. D. G. Gibson et al., Enzymatic assembly of DNA molecules up to several hundred kilobases. Nat. Methods. 6, 343–345 (2009).

36. T. Duong, M. J. Miller, R. Barrangou, M. A. Azcarate-Peril, T. R. Klaenhammer, Construction of vectors for inducible and constitutive gene expression in Lactobacillus. Microb. Biotechnol. 4, 357–367 (2011).

37. M. Teresa Alegre, M. Carmen Rodríguez, J. M. Mesas, Transformation of Lactobacillus plantarum by electroporation with in vitro modified plasmid DNA. FEMS Microbiol. Lett. 241, 73–77 (2004).

38. K. Thompsona ’, M. A. Collinsb, Methods Improveiment in electroporation efficiency for Lactobacillus plantarum biy the inclusion of high concentrations of glycine in the growth medium. J. ofMicrobiological J. Microbiol. Methods. 26, 73–79 (1996).

39. K. Spath, S. Hein, R. Grabherr, “Direct cloning in Lactobacillus plantarum: Electroporation with non-methylated plasmid DNA enhances transformation efficiency and makes shuttle vectors obsolete.” Microb. Cell Fact. 11, 141 (2012).

40. A. a Gomaa et al., Programmable Removal of Bacterial Strains by Use of Genome-Targeting CRISPR/Cas Systems. MBio. 5, e00928–13 (2014).

41. M. Kearse et al., Geneious Basic: An integrated and extendable desktop software platform for the organization and analysis of sequence data. Bioinformatics. 28, 1647–1649 (2012).

42. M. E. Martino et al., Resequencing of the Lactobacillus plantarum strain WJL genome. Genome Announc. (2015), doi:10.1128/genomeA.01382-15.

43. M. Kleerebezem et al., Complete genome sequence of Lactobacillus plantarum WCFS1. Proc. Natl. Acad. Sci. U. S. A. 100, 1990–5 (2003).

44. J. P. Van Pijkeren, R. A. Britton, High efficiency recombineering in lactic acid bacteria. Nucleic Acids Res. 40 (2012), doi:10.1093/nar/gks147.

